# Eco-evolutionary feedbacks shape the evolution of constitutive and inducible defences

**DOI:** 10.1101/2023.04.14.536855

**Authors:** Bridget N. J. Watson, Elizabeth Pursey, Sylvain Gandon, Edze R. Westra

**Affiliations:** ESI, Biosciences, University of Exeter, Cornwall Campus, Penryn TR10 9FE, UK; Centre d’Ecologie Fonctionnelle et Evolutive (CEFE), UMR 5175, CNRS-Université de Montpellier-Université Paul-Valéry Montpellier-EPHE. Montpellier, France

## Abstract

Organisms have evolved a range of constitutive (always active) and inducible (elicited by parasites) defence mechanisms, but we have limited understanding of what drives the evolution of these orthogonal defence strategies. Bacteria and their phages offer a tractable system to study this: bacteria can acquire constitutive resistance by mutation of the phage receptor (surface mutation, *sm*) or induced resistance through their CRISPR-Cas adaptive immune system. Using a combination of theory and experiments we demonstrate that the mechanism that establishes first has a strong advantage because it weakens selection for the alternative resistance mechanism. As a consequence, ecological factors that alter the relative frequencies at which the different resistances are acquired have a strong and lasting impact: high growth conditions promote the evolution of *sm* resistance by increasing the influx of receptor mutation events during the early stages of the epidemic, whereas a high infection risk during this stage of the epidemic promotes the evolution of CRISPR immunity, since it fuels the (infection-dependent) acquisition of CRISPR immunity. This work highlights the strong and lasting impact of the transient evolutionary dynamics during the early stages of an epidemic on the long-term evolution of constitutive and induced defences, which may be leveraged to manipulate phage resistance evolution in clinical and applied settings.

## Introduction

Organisms have evolved a large repertoire of defence systems that offer protection against infectious diseases. Some of these defences are always active – known as constitutive defences – whereas others are elicited by parasites – known as inducible defences (Tollrian and Harvell, 1999). Fitness trade-offs associated with these defences tend to manifest accordingly (i.e. constitutive or infection-induced (Kraaijeveld and Godfray, 1997, Moret and Schmid-Hempel, 2000)), and consequently, organisms are predicted to invest more in constitutive defences as the infection risk increases, and less in induced defences (Westra, et al., 2015). Yet it is not clear what influences the initial evolution of each resistance strategy, or if there are interactions between these alternative strategies that influences their long-term coevolution.

Bacteria and their viruses, called bacteriophages or phages, are a useful model system to study the evolution of different defence strategies. Bacteria can evolve resistance against phages through a wide range of mechanistically distinct defence strategies (Hampton, et al., 2020). Many of those provide innate immunity and are therefore key for determining the levels of pre-existing phage resistance but are less important for the evolution of *de novo* phage resistance (van Houte, et al., 2016). Rapid evolution of phage resistance typically relies on either mutation of the phage receptor, in order to prevent phage adsorption to the bacterial cell, or the acquisition of CRISPR-Cas adaptive immunity, which is based on insertion of phage-derived sequences (spacers) into CRISPR loci in the host genome that are used as a genetic memory to detect and destroy phages during re-infection (Watson, et al., 2021). Evolution of phage resistance by the opportunistic human pathogen *Pseudomonas aeruginosa* PA14 against its obligatory lytic phage, DMS3*vir*, is a case in point, relying either on mutation of the Type IV pilus (surface mutation, *sm*), or its type I-F CRISPR-Cas system. Mutations in the Type IV pilus carry a constitutive cost of resistance, whereas CRISPR immunity is associated with an infection-induced cost (Westra, et al., 2015, Alseth, et al., 2019, Meaden, et al., 2020). Since both mutations confer (almost) perfect resistance to phages, cells that carry these two mutations would not benefit from higher fitness because resistance would not be much higher. In other words, there is strong negative epistasis in fitness between these mutations. This negative epistasis is known to strongly influence the trajectories of adaptation (Day and Gandon, 2012, Hansen, 2013, Bank, 2022). In accordance with theory, selection favours CRISPR immune bacteria over surface mutants when phage densities are low, but the balance tips in favour of surface mutants as phage densities increase (Westra, et al., 2015). In this study, we combine theory and experiments with this bacteria-phage model to explore if and how the short-term transient evolutionary dynamics of these different resistances impacts their long-term evolution.

## Results

To address this gap in our knowledge, we first generated a mathematical model to identify the key parameters that influence the transient evolution of defence mechanisms in an initially sensitive bacterial population when it is exposed to phages (**Fig. 1**). The model allows the joint evolution of the two mechanisms of resistance that can be acquired by susceptible cells via mutation (surface mutation resistance, *sm*) or acquisition of a new spacer (CRISPR resistance) and accounts for possible costs of resistance (fixed cost for surface resistance and conditional cost for CRISPR resistance). This deterministic model allows us to track how initial conditions affect the epidemiology and evolution of the system and thus make predictions on the final frequency of the different types of resistance (more details of the model are presented in **Supplementary Information**).

**Figure 1.**
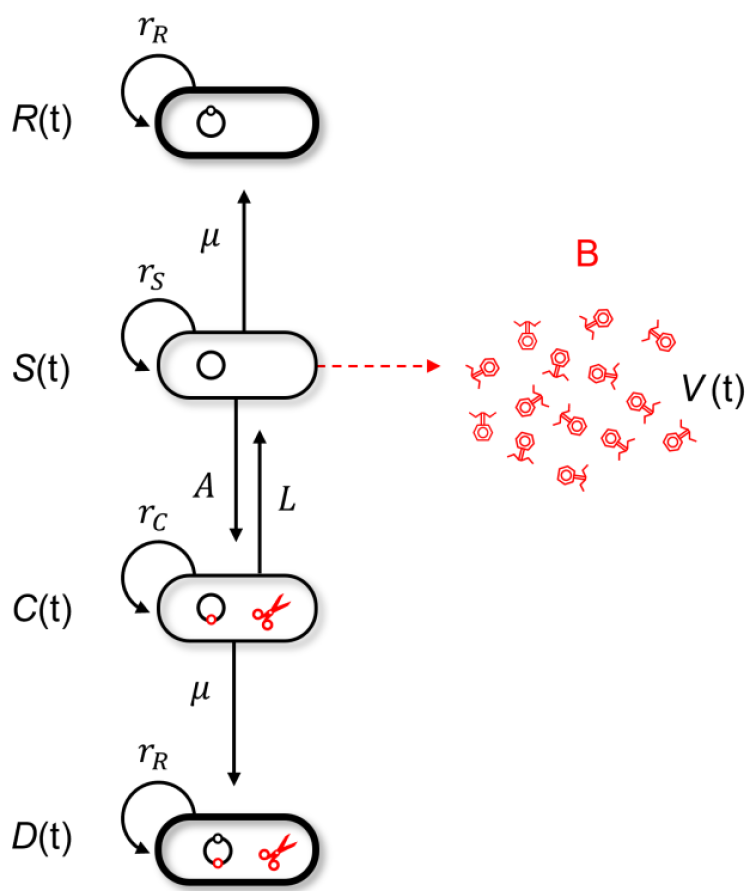
Schematic representation of the model. Naive and uninfected hosts (S hosts) reproduce at rate *r*_*S*_. Upon infection by the phages, they release a burst size *B* of new viral particles. Two distinct types of resistance may emerge: surface modification (*R* hosts) or CRISPR resistance (*C* hosts). *R* hosts reproduce at a rate 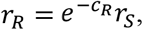 where *c*_*R*_ measures the cost of resistance. *C* hosts reproduce at a rate 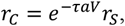 where τ measures the toxicity induced by CRISPR immunity when resistance cells are exposed to the virus. *D* hosts reproduce at a rate 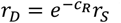 because even though they have both types of resistance, they only express the cost of constitutive resistance because they are never infected by phages. Cells acquire surface modification at rate *μ* (this rate is constant) and acquire CRISPR resistance at rate *AaV* (this rate varies with the exposition to viral particles).

Analysing the change in frequency of the different resistance forms revealed that the initial phase of evolution is key. The more rapidly CRISPR immune bacteria increase in frequency, the more they interfere with selection for surface resistance for two main reasons. First, the increase in resistance to phages in the bacterial population reduces the fitness benefit associated with a new resistance mechanism. Second, the increase in resistance to phages feeds back on viral dynamics and the drop in phage density reduces the selective pressure for *sm*. This causes the spread of this alternative form of resistance to slow down (interference in the selection coefficients is derived in the **Supplementary Information**). Consequently, factors that increase the early acquisition of spacers (relative to the acquisition of mutations in phage receptor genes) will promote the evolution of CRISPR-based immunity, and vice versa for the evolution of surface-based resistance (**Figs. S1 & 2**). Since the acquisition of receptor mutations is tightly linked to bacterial replication, a key prediction from the model is therefore that the amount of replication that can occur in the initially sensitive population until carrying capacity is reached has a major impact on the type of resistance that emerges (**Fig. 2A-C, Fig. S3**). Another prediction from the model is the lack of double resistance (**Figs. S1-4**). Indeed, double resistance has the same level of resistance as single resistance, and it carries the same cost as surface resistance. This implies that epistasis between the two resistance mechanisms is strongly negative. This negative epistasis is expected to yield negative linkage disequilibrium (**Fig. S4**) and a low frequency of double resistance.

**Figure 2.**
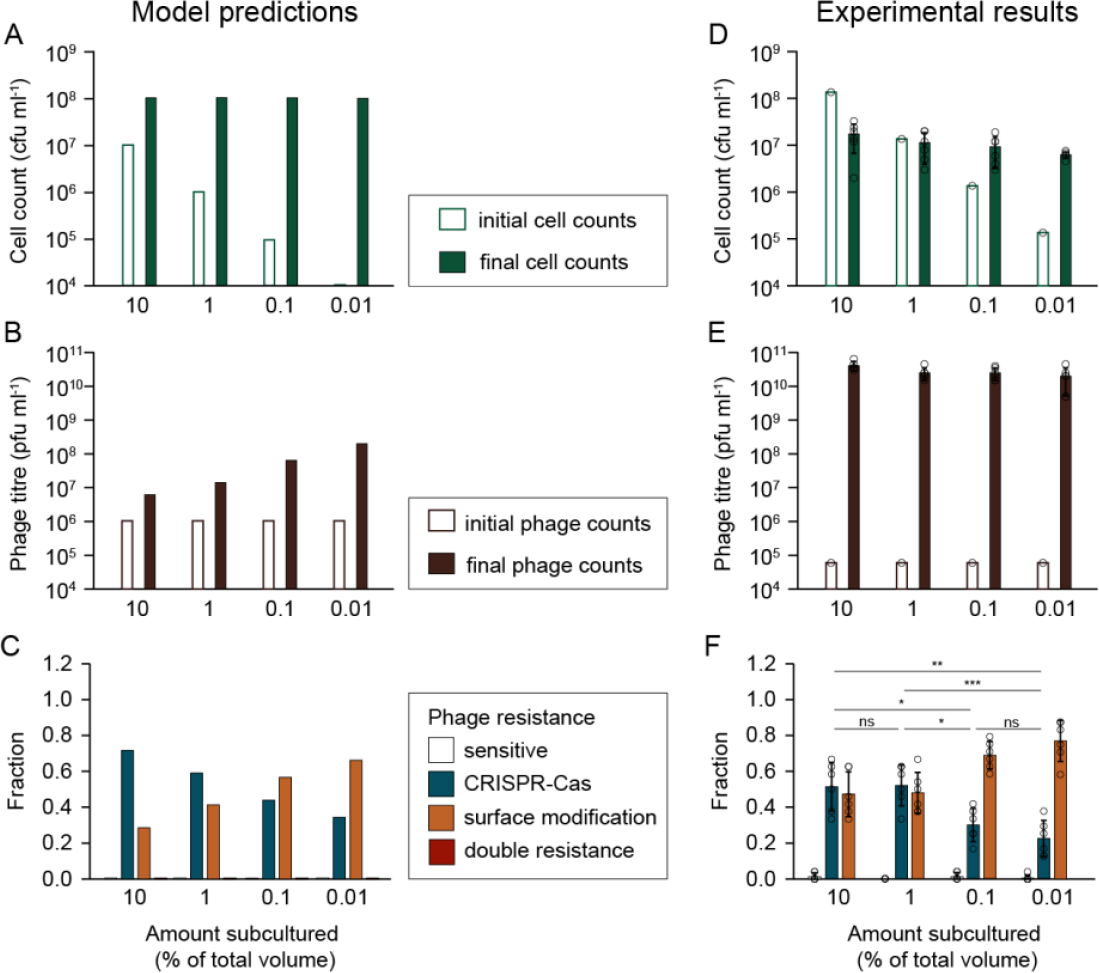
The emergence of surface mutants is replication dependent. Cultures were inoculated with different proportions of stationary phase, sensitive WT *P. aeruginosa* (600 μl: 10%, 60 μl: 1%, 6 μl: 0.1% and 0.6 μl: 0.01% of final volume), and 10^5^ pfu ml^-1^ DMS3*vir* phages. Plots show: (**A, D**) cell counts and (**B, E**) phage counts for each treatment (white bars: initial, green/purple bars: final (1 day)), as well as (**C, F**) the fraction of each resistance type was determined (for 24 clones per replicate, white: phage sensitive, blue: CRISPR-Cas immune, orange: surface-based resistance). (**A-C**) the outcomes predicted by the model (see Supplemental Information) and (**D-F**) show the experimental results. Data shown are the mean ± 1 standard deviation, 6 replicates per treatment. (**F**) Statistical significance between the fractions of CRISPR resistance for each treatment was testing using ANOVA with post-hoc Tukey test, 10% vs 1%: *p* = 0.9995 (ns, not significant), 10% v 0.1%: *p* = 0.0164 (*), 10% v 0.001%: *p* = 0.0011 (**), 1% v 0.1%: *p* = 0.0130 (*), 1% v 0.01%: *p* = 0.0009 (***), 0.1% v 0.01%: *p* = 0.6463 (ns).

Next, we performed evolution experiments with *P. aeruginosa* PA14 and its non-lysogenic phage, DMS3*vir*, to test this model prediction. *P. aeruginosa* PA14 carries a type I-F CRISPR-Cas immune system (Cady, et al., 2012) and is commonly used as a model to study the evolutionary ecology of CRISPR-Cas systems (Westra, et al., 2015, Watson, et al., 2021). We inoculated the same volume of fresh growth media with different amounts of sensitive cells from an overnight culture of WT PA14, ranging from 10% to 0.01% of the total final volume (**Fig. 2**). As a result, the number of rounds of replication until the cultures reached carrying capacity differed between treatments, which in turn affected the opportunity for *sm* clones to emerge in those cultures. Each bacterial culture was also infected with phage DMS3*vir* (10^5^ pfu ml^-1^). Despite the differences in initial cell concentrations, all cultures reached similar final counts (**Fig. 2D**) as did the phage counts (**Fig. 2E**). Crucially, as predicted by the model (**Fig. 2A-C, Fig. S3**), cultures with the smallest inoculum of 0.01% mostly evolved surface-based resistance (*sm* fraction: 0.77± 0.11), whereas those with the largest inoculum of 10% mostly evolved CRISPR immunity (CRISPR fraction: 0.51± 0.14) (**Fig. 2F**). The proportion of CRISPR immunity that evolved significantly increased with the bacterial inoculum size (ANOVA, *p* = 0.0002, with post hoc Tukey test, 10% vs 1%: *p* = 0.9995, 10% v 0.1%: *p* = 0.0164, 10% v 0.001%: *p* = 0.0011, 1% v 0.1%: *p* = 0.0130, 1% v 0.01%: *p* = 0.0009, 0.1% v 0.01%: *p* = 0.6463). These data therefore show that higher levels of CRISPR immunity emerged when less bacterial replication occurred.

Our model also predicts that increasing phage infection will promote the evolution of CRISPR-Cas immunity. This is because CRISPR immunity only evolves following phage infection whereas surface mutants emerge during replication; increasing the initial phage dose thus increases the early acquisition of CRISPR immunity relative to the acquisition of surface resistance. Likewise, when bacteria grow to higher population densities, phages will reach higher densities, more rapidly, as they will be more likely to find a host when they diffuse through the media (**Fig. S1**). Note, however, that because CRISPR immunity is associated with an infection-induced toxicity cost (Westra, et al., 2015), selection favours bacteria with surface resistance when the density of viruses becomes too high (without toxicity cost, CRISPR immune bacteria are always favoured at higher viral doses, see **Fig. S2**).

To test these model predictions, we experimentally examined, using a full factorial design, the effect of phage exposure by varying the initial doses (10^1^, 10^2^, 10^3^, 10^5^ and 10^9^ plaque forming units (pfus) ml^-1^) and the carrying capacity of the media (0.0002, 0.002, 0.02 and 0.2% glucose, **Fig S5A**) in infection experiments of PA14 with DMS3*vir*. We analysed the cell and phage densities across the experiment and saw that both conditions had a significant impact on the phage densities at t=1-, 2- and 3-days post infection (dpi), as predicted (**Fig. S5A&B**). In conditions with low phage doses and/or low carrying capacity (low glucose concentration), phage counts increased over the duration of the experiment (**Fig S5C**). Whereas in conditions with high phage doses and/or high carrying capacities, phage densities peaked with high counts by t=2 dpi (**Fig. S5C**). Resistance profiles were then determined daily for three days to capture the dynamics of resistance evolution across these treatments. Visual inspection of the resulting data supported the model prediction: generally, populations appear to evolve higher levels of CRISPR immunity when either the initial phage dose or the carrying capacity is high (**Fig. 3A**). For example, following infection with 10^1^ pfu of DMS3*vir* (in high glucose conditions), CRISPR immunity was barely detected at 1 dpi (CRISPR fraction 10^1^: 0.09± 0.10, mean± 1 standard deviation) but with doses of 10^3^ pfu and higher, most clones had CRISPR immunity at 1 dpi (CRISPR fraction 10^3^: 0.59± 0.14 10^5^: 0.70± 0.10, 10^9^: 0.64± 0.02) (**Fig. 3A, Fig. S6A**). Similarly, for the carrying capacity conditions, in the treatments with low carrying capacity (0.0002 and 0.002% glucose), CRISPR clones were not detected 1 dpi, but with high carrying capacity, CRISPR immunity was dominant (CRISPR fraction 0.02% glucose: 0.49± 0.25, 0.2% glucose: 0.59± 0.14) (**Fig. S6B**). To analyse these data more rigorously, we developed statistical models. We used separate mixed effects models to assess the relative contributions of each variable (glucose concentration, initial phage dose, final phage density and final cell density) for each timepoint, controlling for treatment replicate. Model selection was then used to determine which variables were most important for explaining CRISPR evolution at each timepoint (**Fig. 3B-D**).

**Figure 3.**
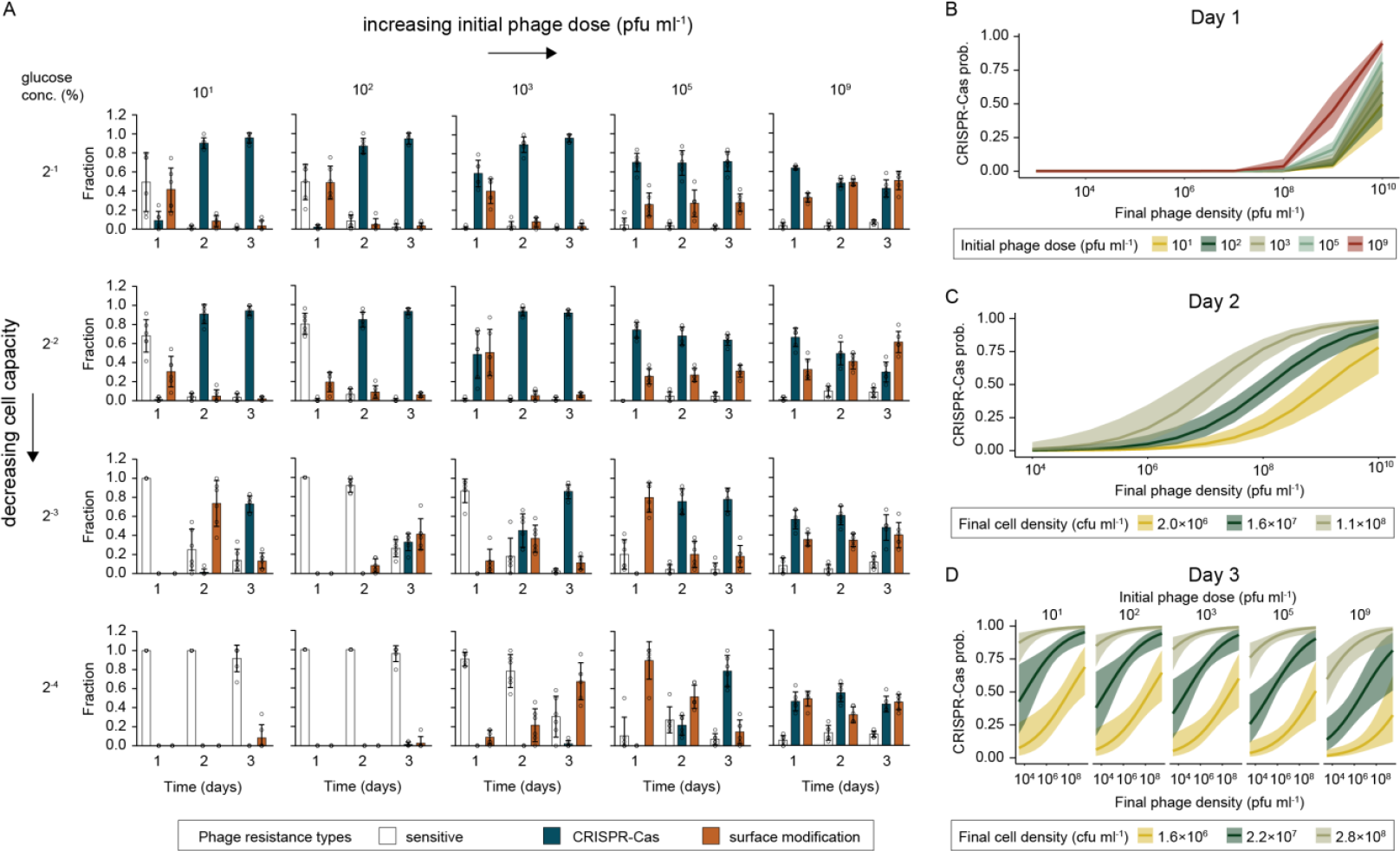
Higher levels of CRISPR immunity are observed with higher phage exposure. (**A**) Fraction of each resistance type (white: phage sensitive, blue: CRISPR-Cas immune, orange: surface-based resistance) over three days of evolution following exposure of initially phage sensitive WT *P. aeruginosa* to different amounts of DMS3*vir* phages (10^1^, 10^3^, 10^5^, 10^9^ pfu ml^-1^) and in media containing different levels of glucose (0.2, 0.02, 0.002, 0.0002% glucose, resulting in different carrying capacities, see Fig. S4A). Data shown are the mean ± 1 standard deviation, 6 replicates per treatment, 24 clones tested per replicate. (**B-D**) Prediction plots showing model-estimated means and 95% confidence intervals based on statistical modelling of the data (in **A**), in which model selection was used to retain the most important predictors of CRISPR evolution on (**B**) Day 1, (**C**) Day 2 and (**D**) Day 3.

At 1 dpi, final phage density and the initial phage dose were the most important variables explaining the probability that a clone evolves CRISPR, with more CRISPR evolution predicted as these variables increased (**Fig. 3B**). At 2 dpi, final phage density and final cell density explained the most variation, with higher phage and cell densities both predicting an increased probability of CRISPR immunity (**Fig. 3C**). Finally, by 3 dpi, initial phage dose, final phage density and final cell density were all retained in the model (**Fig. 3D**). In this case, higher phage doses at the start of the experiment were associated with a slightly reduced probability of CRISPR evolution by 3 dpi. Notably, at this final timepoint, measured final phage and cell densities had far larger impacts on CRISPR evolution than the initial phage inoculum, and CRISPR evolution was predicted at substantially lower phage and cell densities than at 2 dpi. Glucose concentration was not retained in any model, suggesting that its impact on cell density was indeed the main driver of the effects seen. Collectively, these results indicate that conditions of high phage exposure, be it due to high dose of phage or high carrying capacity, are associated with higher levels of CRISPR immunity.

## Discussion

Here we combined novel theory and experimental work to examine the transient evolutionary dynamics of induced and constitutive defences. Using *P. aeruginosa* and phage DMS3*vir* as a model system, we examined the factors driving the relative abundance of CRISPR immunity (induced) and surface mutants (*sm*, constitutive) that emerge in phage sensitive bacterial populations. The joint evolution of induced and constitutive defences has previously been modelled, using an adaptive dynamics framework to identify the evolutionary stable investment in the two strategies (Westra, et al., 2015). This analysis showed the influence of the exposure to phage predation and the productivity of the environment on the long-term coevolutionary outcomes between these two alternative resistance strategies. Here, we develop a model where we compete different bacterial genotypes that either carry or lack the resistance at each of two resistance loci, resulting in four distinct bacterial genotypes. This model allows us to track the transient dynamics of different resistance genotypes in combination with phage density. Our model shows that the emergence of the first resistance mechanism interferes with the evolution of the alternative resistance and may thus affect the long-term evolutionary outcome.

We see that *sm* arise independent of phage infection but during DNA replication and we predicted, and experimentally demonstrated, that *sm* relative abundance in the population is dependent on the replication potential of the populations. Our finding that bacteria increasingly rely on their CRISPR-Cas immune systems under conditions of low bacterial growth is consistent with previous studies showing that CRISPR-Cas immune systems become relatively more important when the focal species is cultured in resource-limited growth media (Westra, et al., 2015), when they are exposed to bacteriostatic antibiotics (Dimitriu, et al., 2022) or when they compete with other bacteria (Alseth, et al., 2019). On the other hand, CRISPR immunity evolution is dependent on phage infection. Hence, increasing the cell culture carrying capacity and the number of phages initially present resulted in faster phage epidemics and hence, greater phage exposure. The finding that higher phage densities promote evolution of CRISPR immunity is consistent with the positive correlation between CRISPR and phage prevalence in metagenome sequence data (Meaden, et al., 2022). However, even though high phage densities fuel the rate at which CRISPR immunity is acquired, bacteria that evolved *sm* resistance will dominate at very high phage densities, due to the compounding costs associated with CRISPR immunity that are induced by infection (Westra, et al., 2015, Meaden, et al., 2020). Indeed, our statistical model and experimental data support the notion that if phage titres are high following the emergence of resistance types, selection favours the invasion of *sm* resistance; for example, at the highest phage exposure treatments bacterial cultures grown with 0.2% or 0.02% glucose contained approximately equal proportions of CRISPR immune bacteria and surface mutants, whereas cultures were dominated by CRISPR immune bacteria in the treatments with lower phage exposures.

Whichever of these two different resistance mechanisms, CRISPR-Cas or *sm*, dominates the evolutionary response against phages has major implications for the ability of this pathogen to cause disease. Specifically, evolution of *sm* resistance is associated with large virulence trade-offs that are not detected when the bacteria evolve CRISPR-based immunity (Alseth, et al., 2019). Understanding when CRISPR-Cas immunity and surface-based resistance evolve is therefore critical, especially since empirical studies support the idea that therapeutic application of phages is much more effective when virulence trade-offs are triggered in resistant bacteria (Kortright, et al., 2019).

## Acknowledgements

This work was funded by a grant from the Natural Environment Research Council (NE/S001921/1) and a grant from the European Research Council (https://erc.europa.eu) (ERC-STG-2016-714478 - EVOIMMECH) awarded to E.R.W. B.N.J.W acknowledges support from the Biotechnology and Biological Sciences Research Council (BB/X010600/1).

## Methods

### Mathematical model

See **Supplementary Information** and **Figures S1-4**.

### Bacterial strains and phage

Bacterial strains used in this study include *Pseudomonas aeruginosa* UCBPP-PA14 (WT), UCBPP-PA14 *csy3::lacZ* (KO) (Cady, et al., 2012), UCBPP-PA14 BIM2 with two spacers targeting DMS3*vir* (BIM) (Westra, et al., 2015) and UCBPP-PA14 *csy3::lacZ* spontaneous surface mutant (*sm*) (Westra, et al., 2015). Phages used in this study include the obligatory lytic temperate phage, DMS3*vir* (Cady, et al., 2012), and DMS3*vir* carrying anti-CRISPR (Acr) IF1 (Landsberger, et al., 2018).

### Experimental evolution

Evolution experiments with PA14 (WT) and DMS3*vir* were performed as previously (Westra, et al., 2015). Briefly, 6 ml cultures of M9 containing 0.2, 0.02, 0.002 or 0.0002% glucose in glass vials (*n* =6) were inoculated with approximately 10^6^ colony forming units (cfus) of PA14 (1:1000 subculture of M9 (0.2% glucose) adapted cells). Changing the glucose concentration changes the carrying capacity of the media but growth rate is unaffected (**Fig. S5A**). Phages were added to each vial in varying amounts (10^1^, 10^2^, 10^3^, 10^5^, 10^9^ plaque forming units (pfu)), before the vial lids were tightly closed and cultures were incubated at 37 degrees with shaking. Cultures were subcultured 1:100 daily, for 3 days, and cfu and pfu counts were determined daily by plating and spot assays (**Fig. S5B&C**). To determine the phage resistance phenotypes, 24 clones were randomly selected from each replicate, inoculated into LB in a 96-well plate and grown overnight. Cultures were streaked against phages DMS3*vir* and DMS3*vir-*AcrIF1. Consistent with previous work (Westra, et al., 2015, Rollie, et al., 2020, Chevallereau, et al., 2020), phage sensitive clones were susceptible to both phages, *sm* were resistant to both phages and the CRISPR clones were resistant to DMS3*vir*, but not DMS3*vir*-AcrIF1. To test the effect of different inoculum amounts, varying amounts of 6 ml M9 (0.2% glucose) cultures were subcultured (600 μl: 10%, 60 μl: 1%, 6 μl: 0.1% and 0.6 μl: 0.01%) into fresh media, for a total final volume of 6 ml. 10^5^ pfus were added to each culture (*n* =6) and phage resistance profiles were determined following 1 day of growth.

### Statistical modelling

Mixed effects models were constructed to examine the relative contributions of all potential predictors on CRISPR evolution. A binomial dataset was constructed where CRISPR evolution was coded as 1 or 0 for each clone per replicate. Next, a binomial generalised linear mixed effects model with fixed effects of glucose, cell density, phage density and initial phage inoculum, and treatment replicate as a random effect, was run for each timepoint (day 1-3). Cell and phage density data were log transformed. A maximal model was generated and all possible candidate models were compared using the AIC method with *dredge* from the MuMIn package (Barton, 2009). AIC values assess the fit of a model by looking at the likelihood of a model given the data, penalising for increased number of parameters (as increased complexity of the model increases parameter uncertainty). We selected the most parsimonious model (i.e. the model with fewest parameters) within 2 delta AICs for each timepoint. Model comparisons based on AIC are presented in **Table S1**, with the selected models highlighted. Model estimates for the selected models are presented in **Table S2**.

For the selected models, prediction data frames with 95% confidence intervals were generated using *ggpredict* from the package ggeffects (Lüdecke, 2018), model dispersion was tested and scaled residuals were examined using DHARMa residual diagnostics (Hartig, 2020), and the final predictions were visualised with ggplot2 v3.3.2 and the wesanderson package. All statistical analyses were performed in R v4.0.2, and code is available at https://github.com/elliekpursey/Watson_2022.

## Supporting Information

### Transient selection for different types of resistance against bacteriophages

#### 1. The model

We want to model the transient evolution of a bacterial population exposed to phage predation. Initially, the bacteria population is fully susceptible (*S* is the density of susceptible, or sensitive, bacteria) and is exposed to a lytic phage (*V* is the density of free viral (phage) particles). Susceptible bacteria can either evolve surface resistance by mutation (*R* is the density of surface resistant bacteria) or they can evolve CRISPR resistance after the acquisition of a new spacer targeting the phage (*C* is the density of CRISPR resistant bacteria). Note that CRISPR resistant cells may also acquire surface resistance by mutation (*D* is the density of cells that carry both surface resistance and CRISPR resistance) (Figure 1). The following dynamical system allows us to track the transient build-up of resistance via these two different mechanisms:

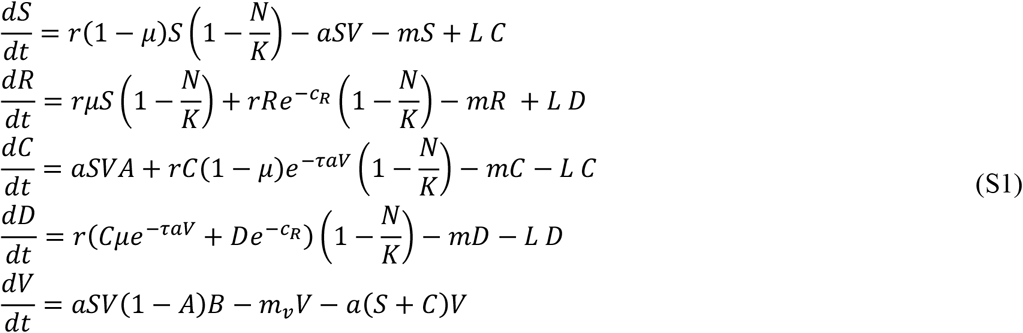

where we assume that surface resistance carries a fitness cost *c*_*R*_ while CRISPR resistance may suffer from a fitness cost τ when the CRISPR resistant cells are infected by the phage (this toxicity increases with the force of infection *aV*).

The total bacteria population size is defined as: *N* = *S* + *R* + *C* + *D*. The change in the total population size is given by:

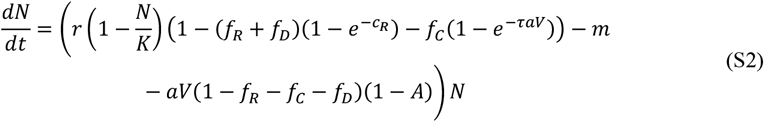

where the frequencies of the different types of resistant cells are indicated as:

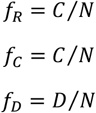

#### 2. Evolution of resistance

Next, to understand the transient evolutionary dynamics of resistance, we focus on the dynamics of the frequencies of the different types of resistance.

##### 2.1 Evolution of surface resistance

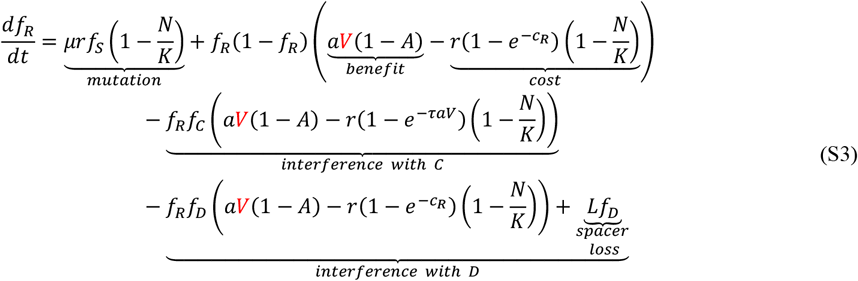

The first term is the flux of mutation from susceptible bacteria with *f*_*S*_ = *S*/*N*.

The second term depends on the amount of genetic variation (i.e. *f*_R_(1 − *f*_R_)) and the strength of selection on surface resistance (the benefit minus the cost): (i) the benefit is to escape from phage predation, (ii) the cost depends on bacteria density because reproduction is density dependent.

The third term captures the potential interference with CRISPR resistance. It is usually negative (the presence of an alternative form of resistance reduces the competitive edge of this mutation) but this effect is reduced by the cost on CRISPR resistance.

The final two terms capture the potential interference with double resistant cells. It is usually negative (the presence of an alternative form of resistance reduces the competitive edge of this mutation) but this negative effect is reduced by the cost on CRISPR resistance, as well as the loss of spacer(s) from the double resistant cells.

##### 2.2 Evolution of CRISPR resistance

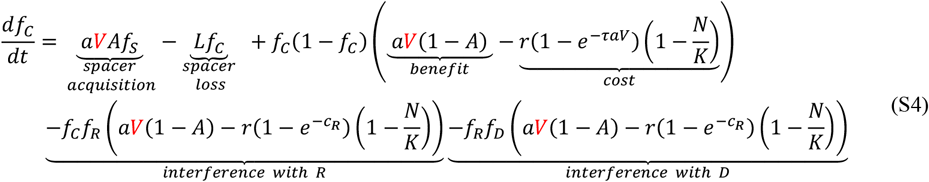

The first two terms correspond to the rate of spacer acquisition (influx of CRISPR resistance) and spacer loss (outflux of CRISPR resistance). In contrast to surface resistance, the influx of CRISPR resistance depends on viral density and does not depend on bacteria reproduction.

The second term depends on the amount of genetic variation (i.e. *f*_*C*_ (1 − *f*_*C*_)) and the strength of selection on CRISPR resistance (the benefit minus the cost): (i) the benefit is to escape from phage predation, (ii) the cost depends on bacteria density because reproduction is density dependent but it also depends on phage density (because the cost of CRISPR resistance is only expressed when bacteria are infected).

The third term captures the interference with the surface resistance. It is usually negative (the presence of an alternative form of resistance reduces the competitive edge of this mutation) but this effect is reduced by the cost of surface resistance.

The fourth term captures the interference with the double resistance cells. It is usually negative (the presence of an alternative form of resistance reduces the competitive edge of this mutation) but this effect is reduced by the cost on surface resistance (double resistance cells do not suffer from the toxicity cost because the phage does not enter the cells).

##### 2.3 Evolution of double resistance

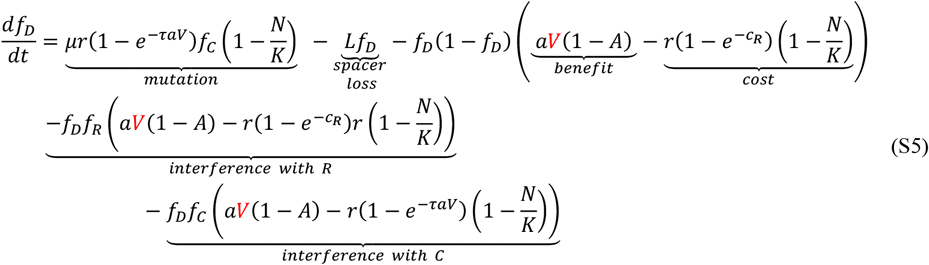

The first term is the flux of mutation from CRISPR resistant bacteria that acquire a surface resistance.

The second term results from the loss of the CRISPR spacer(s) from the double resistant cells.

The third term captures the potential interference with surface resistant cells and depends on the amount of genetic variation (i.e. *f*_*D*_(1 − *f*_*D*_)) and the strength of selection on resistance (the benefit minus the cost): (i) the benefit is to escape from phage predation, (ii) the cost depends on bacteria density because reproduction is density dependent. It is usually negative (the presence of an alternative form of resistance reduces the competitive edge of this mutation) but this effect is reduced by the cost on surface resistance.

The fourth term captures the potential interference with CRISPR resistant cells. It is usually negative (the presence of an alternative form of resistance reduces the competitive edge of this mutation) but this negative effect is reduced by the cost on CRISPR resistance.

## 3 Figures

**Figures S1** and **S2** show numerical simulations that mimic batch transfer experiments when one varies the initial doses of viruses and the carrying capacity of the bacterial populations. The comparison between (S1) and (S2) can be used to make specific predictions on the transient evolution of the different forms of resistance. The influx of surface resistance is governed by the mutation rate μ and the rate of reproduction. The influx of CRISPR resistance is governed by the acquisition rate *A* and the density of viruses. Higher initial doses of free viruses are going to promote the acquisition of spacers and hence promote the evolution of CRISPR (versus surface resistance). Other parameters can also act on viral replication (like carrying capacity) and favour the evolution of CRISPR.

**Figure S3** shows the outcome under different initial conditions to understand the effect of host replication on resistance evolution.

**Figure S4** shows the transitory dynamics of the densities of the different types of cells and the density of free viruses, as well as the dynamics of the linkage disequilibrium between the two resistance loci. This linkage disequilibrium is always negative because the accumulation of different resistance mechanisms does not increase the amount of resistance (negative epistasis).

**Figure S1.**
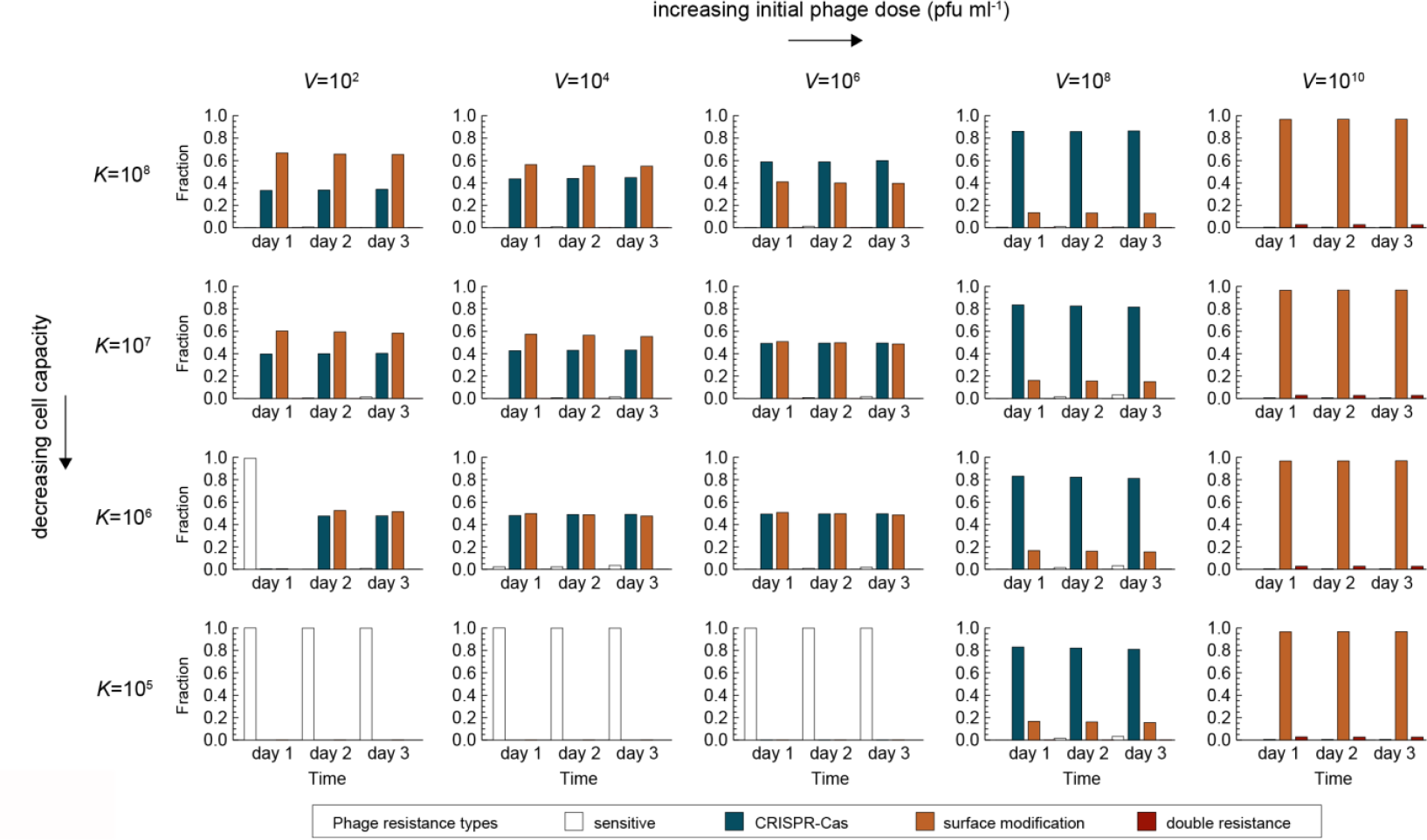
Emergence of phage resistance over time. Frequency of the different bacterial resistance types (*S* (sensitive, white bars), *C* (CRISPR immune, blue), *R* (surface mutants, orange) or *D* (both CRISPR and surface mutants, red)) through time (3 days) and for different values of: the initial doses of free viruses (*V*), and the carrying capacity (*K*). All the simulations started with an initial density of susceptible cells at *K*/100. Other parameter values: *r* = 1, *m* = 0, *m*_*v*_= 0, *a* = 10^−8^, *B* = 100, *c*_*R*_ = 0.01, τ = 0.01, *μ* = 10^−4^, *A* = 5 10^−4^, *L* = 10^−3^.

**Figure S2.**
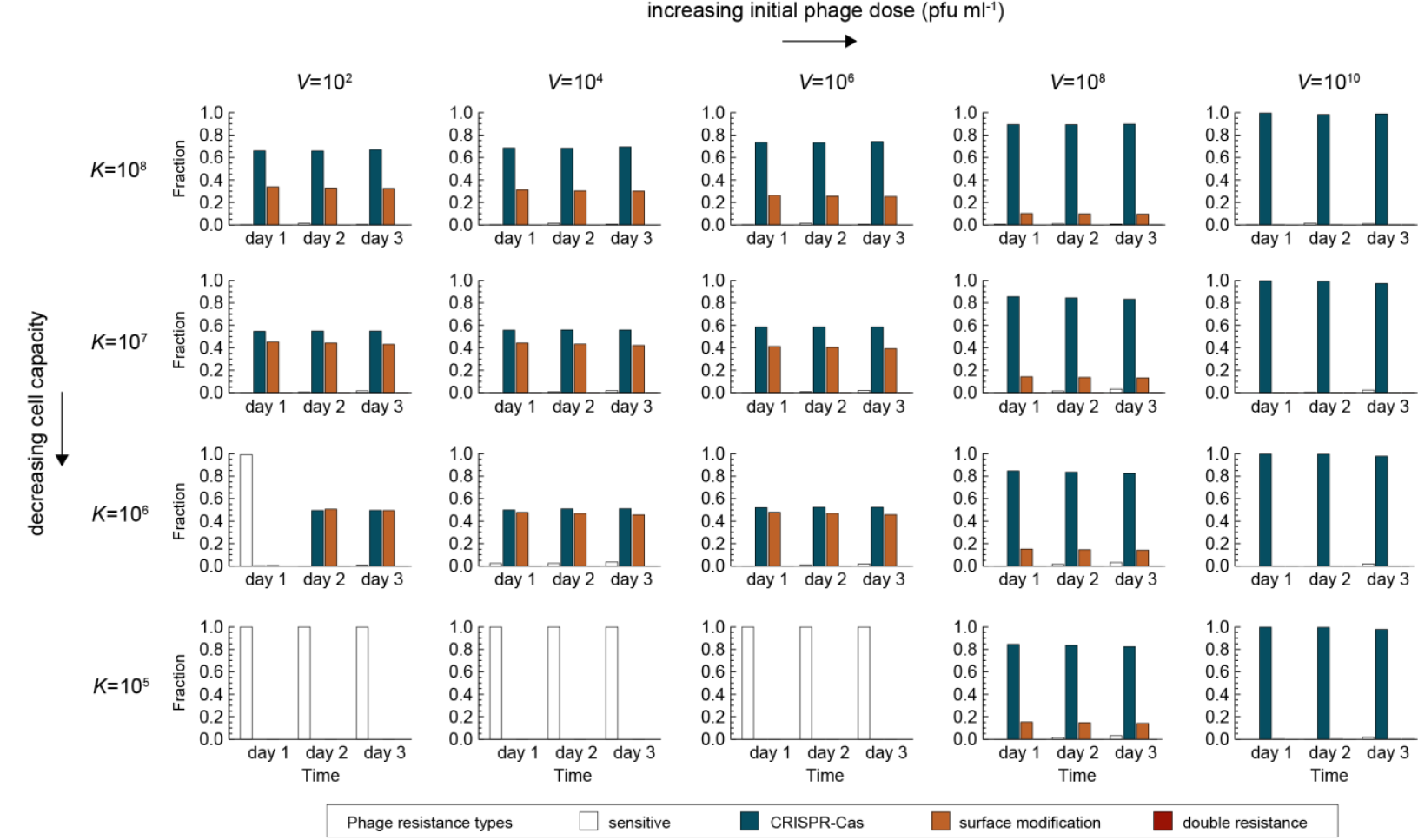
Emergence of resistance over time in the absence CRISPR toxicity. Frequency of the different bacterial resistance types (*S* (sensitive, white bars), *C* (CRISPR immune, blue), *R* (surface mutants, orange) or *D* (both CRISPR and surface mutants, red)) through time (3 days) and for different values of: the initial doses of free viruses (*V*), and the carrying capacity (*K*), in the absence of an induced CRISPR immunity toxicity (τ = 0.0). All the simulations started with an initial density of susceptible cells at *K*/100. Other parameter values: *r* = 1, *m* = 0, *m*_*v*_ = 0, *a* = 10^−8^, *B* = 100, *c*_*R*_ = 0.01, τ = 0.0, *μ* = 10^−4^, *A* = 5 10^−4^, *L* = 10^−3^.

**Figure S3.**
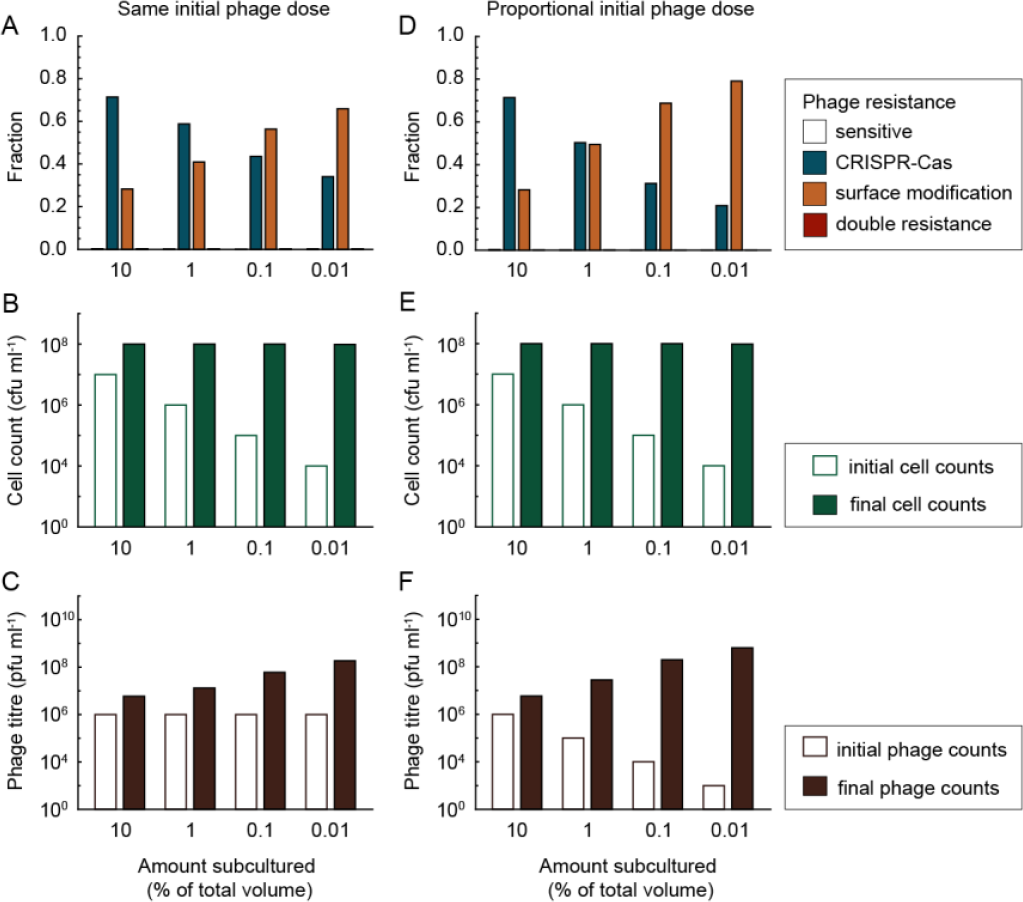
Effect of varying the initial densities of susceptible cells. Plots show: (**A, D**) the predicted resistance type fractions, (**B, E**) cell counts and (**C, F**) phage counts, with (**A-C**) the same initial phage dose (*V* = 10^6^) or (**D-F**) when the initial dose of viruses was diluted to keep the same initial multiplicity of infection of 0.1. Amount of starting inoculum: 10% (*K*/10), 1% (*K*/100), 0.1% (*K*/1000), 0.01% (*K*/10000) of total final volume. Other parameter values: *r* = 1, *m* = 0, *m*_*v*_ = 0, *a* = 10^−8^, *B* = 100, *c*_*R*_ = 0.01, τ = 0.01, *μ* = 10^−4^, *A* = 5 10^−4^, *L* = 10^−3^. Panels (**A-C**) are also shown in Figure 2, alongside the experimental results (Fig. 2D-F).

**Figure S4.**
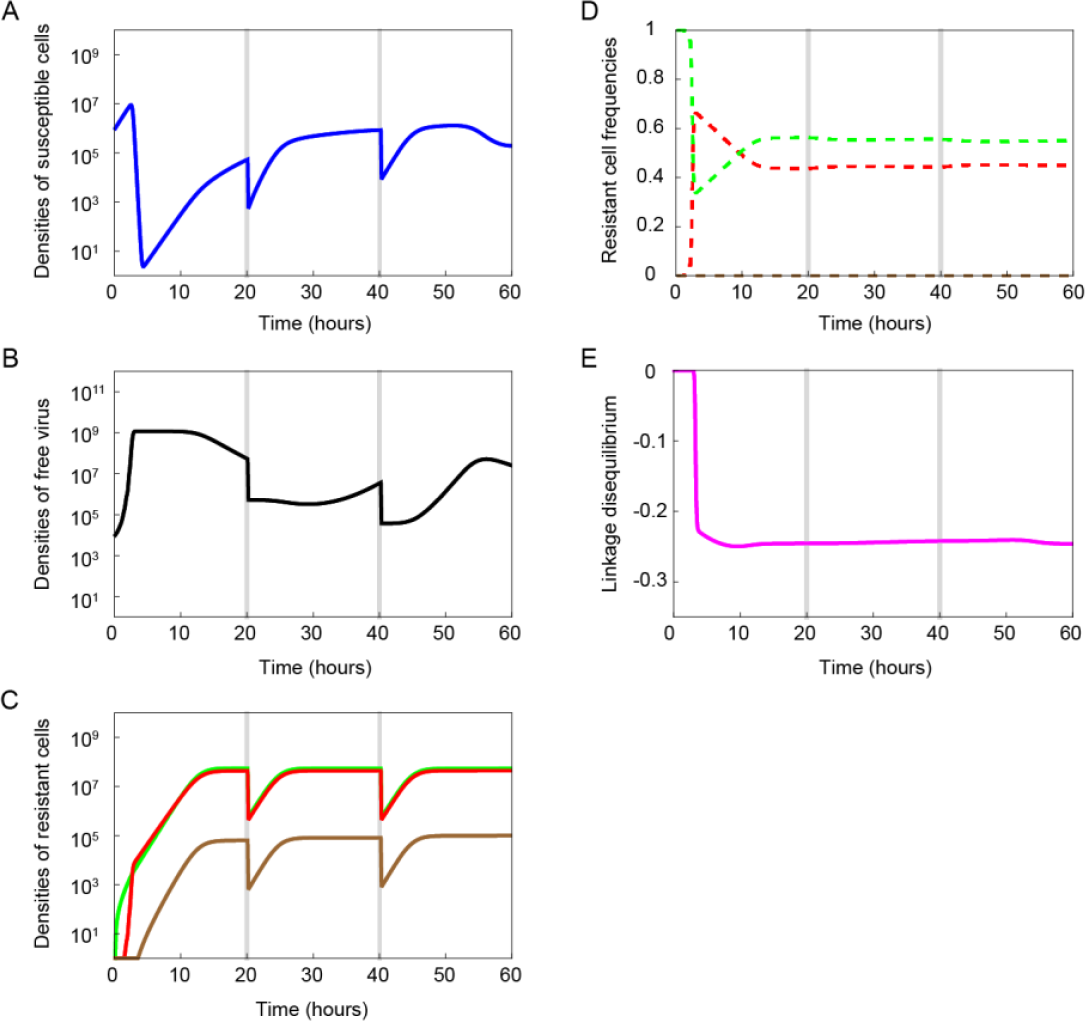
Transitory dynamics of bacteria and viruses across time. The plots show the dynamics for 3 transfers (20 hours between each transfer, indicated by a vertical grey line), including, (**A**) density of *S* cells (susceptible, blue line), (**B**) density of *V* (viruses, black line), (**C**) densities of the different types of resistant cells (*R* (surface mutation, green line), *C* (CRISPR immune, red line) and *D* (both resistances, brown line), (**D**) frequencies of R, *C* and *D* cells (dashed lines with same colours as in panel C) and (**E**) dynamics of linkage disequilibrium between the resistance loci across time. Linkage disequilibrium between the resistance loci is measured as: *LD* = *f*_*S*_*f*_*D*_ − *f*_R_*f*_*C*_. Parameter values: *r* = 1, *m* = 0, *m*_*v*_ = 0, *a* = 10^−8^, *B* = 100, *c*_*R*_ = 0.01, τ = 0.01, *μ* = 10^−4^, *A* = 5 10^−4^, *L* = 10^−3^, *K* = 10^8^ and (initial) *V* = 10^4^.

**Figure S5.**
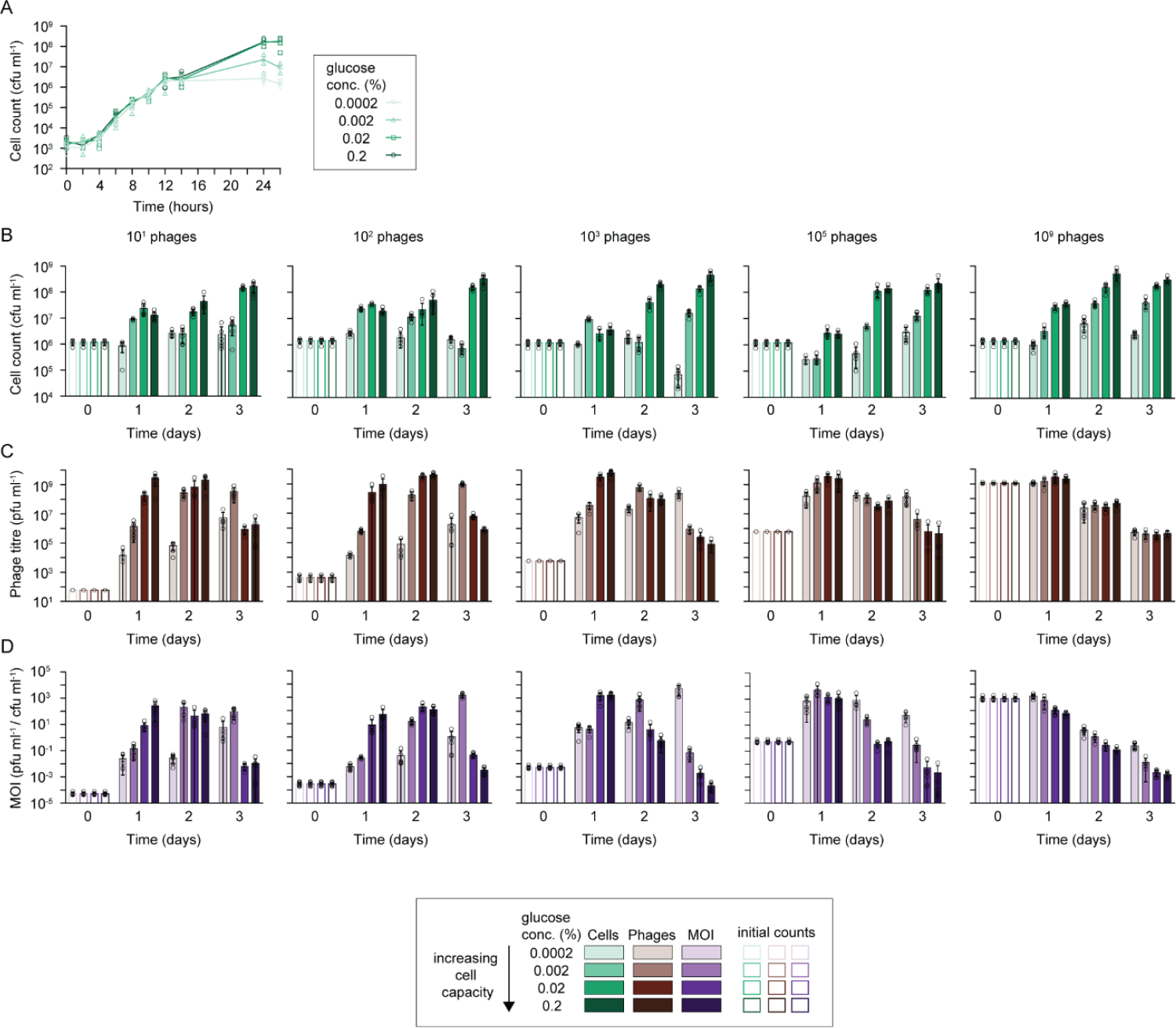
Cell and phage counts from evolution experiment with a range of phage doses and glucose treatments (Figure 3). Plots show (**A**) growth curves of WT cultures grown in M9 containing different amounts of glucose (0.2, 0.02, 0.002 and 0.0002%), (**B**) cell counts (cfu ml^-1^), (**C**) phage counts (pfu ml^-1^) and (**D**) the multiplicities of infection (MOI: phage count/cell count) across the three-day experiment. Initial (day 0) values are depicted as white bars, increasing colour density (light-dark shades) represent increasing glucose concentrations in the growth media/culture carrying capacity.

**Figure S6.**
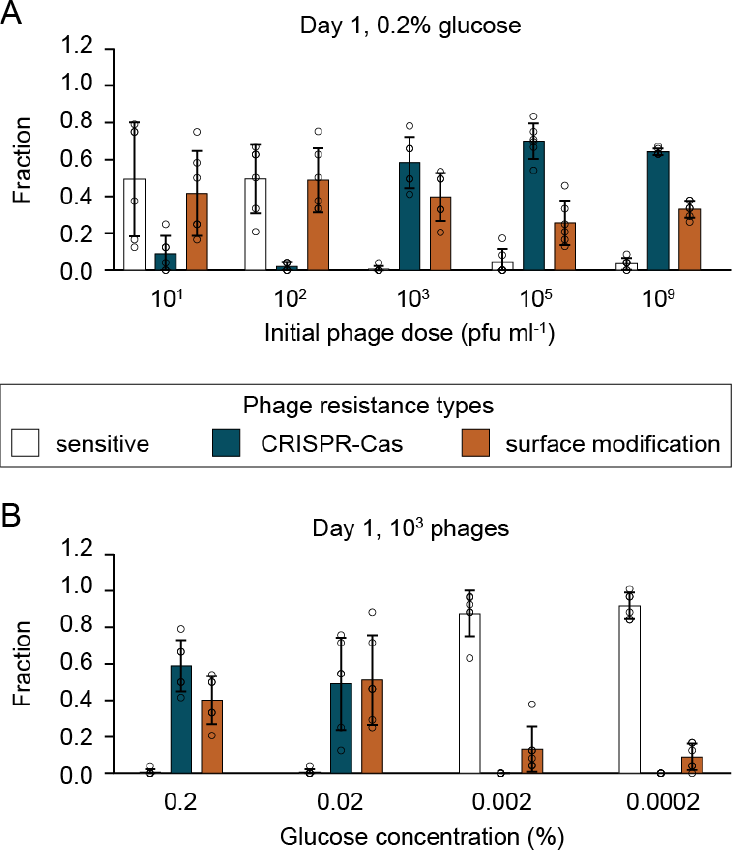
Higher levels of CRISPR immunity are observed with higher phage exposure (selection of data from Fig. 3A). Fraction of each resistance type (white: phage sensitive, blue: CRISPR-Cas immune, orange: surface-based resistance) that evolved after one day of evolution following exposure of initially phage sensitive WT *P. aeruginosa* to: (**A**) different amounts of DMS3*vir* phages (10^1^, 10^3^, 10^5^, 10^9^ pfu ml^-1^) in media containing 0.2% glucose (highest carrying capacity), and (**B**) a moderate phage dose (10^3^ pfu ml^-1^) in media containing different levels of glucose (0.2, 0.02, 0.002, 0.0002% glucose, resulting in different carrying capacities, see Fig. S4A). Data shown are the mean ± 1 standard deviation, 6 replicates per treatment, 24 clones tested per replicate.

**Table S1.** AIC selection tables for binomial generalised linear mixed effects models. Fixed effects of glucose, cell density, phage density and initial phage inoculum, and treatment replicate as a random effect, are shown for each timepoint. Rows with selected models are shown in bold. ‘Phage’ refers to initial phage inoculum size, whilst ‘log_phage’ and ‘log_cell’ indicate measured phage and cell densities, respectively.

| (Intercept) | glucose | log_cell | log_phage | phage | df | logLik | AICc | delta | weight |
| --- | --- | --- | --- | --- | --- | --- | --- | --- | --- |
| **T= 1-day post infection** |  |  |  |  |  |  |  |  |  |
| **-31.4088** | - | - | **3.102901** | **0.364453** | **4** | **-884.912** | **1777.837** | **0** | **0.38751** |
| -30.4731 | 2.333508 | - | 2.966241 | 0.387754 | 5 | -884.182 | 1778.385 | 0.547847 | 0.294659 |
| -32.7203 | - | 0.218796 | 3.087841 | 0.363516 | 5 | -884.555 | 1779.13 | 1.293173 | 0.20299 |
| -31.1974 | 2.00583 | 0.100957 | 2.977234 | 0.383943 | 6 | -884.12 | 1780.269 | 2.43239 | 0.114841 |
| -34.0178 | - | - | 3.585071 | - | 3 | -902.726 | 1811.46 | 33.62259 | 1.94E-08 |
| -34.7215 | -2.15011 | - | 3.682213 | - | 4 | -902.322 | 1812.657 | 34.82042 | 1.06E-08 |
| -35.8714 | - | 0.259674 | 3.597894 | - | 4 | -902.422 | 1812.859 | 35.02156 | 9.63E-09 |
| -38.283 | -3.38692 | 0.439741 | 3.761776 | - | 5 | -901.565 | 1813.151 | 35.31414 | 8.32E-09 |
| -7.29194 | 16.11358 | - | - | 0.784458 | 4 | -926.018 | 1860.05 | 82.21321 | 5.44E-19 |
| -6.1196 | 16.51069 | -0.1785 | - | 0.783321 | 5 | -925.928 | 1861.878 | 84.04071 | 2.18E-19 |
| -11.0008 | - | 0.671885 | - | 0.803605 | 4 | -940.374 | 1888.761 | 110.9244 | 3.17E-25 |
| -6.51748 | - | - | - | 0.800267 | 3 | -941.5 | 1889.007 | 111.1705 | 2.80E-25 |
| -5.0236 | 20.21595 | - | - | - | 3 | -961.545 | 1929.097 | 151.2604 | 5.53E-34 |
| -4.10364 | 20.50837 | -0.13997 | - | - | 4 | -961.522 | 1931.057 | 153.2202 | 2.07E-34 |
| -3.65631 | - | - | - | - | 2 | -971.966 | 1947.937 | 170.1 | 4.48E-38 |
| -9.33632 | - | 0.841333 | - | - | 3 | -971.098 | 1948.204 | 170.3674 | 3.92E-38 |
| **T= 2-day post infection** |  |  |  |  |  |  |  |  |  |
| -23.9903 | - | 1.450115 | 1.599456 | 0.149782 | 5 | -1236.26 | 2482.543 | 0 | 0.49582 |
| **-22.6156** | - | **1.582378** | **1.391144** | - | **4** | **-1238.1** | **2484.206** | **1.663467** | **0.215828** |
| -24.5147 | -0.9059 | 1.512292 | 1.617788 | 0.145486 | 6 | -1236.22 | 2484.463 | 1.919925 | 0.189853 |
| -23.7645 | -1.96563 | 1.70658 | 1.437049 | - | 5 | -1237.88 | 2485.775 | 3.232544 | 0.098489 |
| -14.6326 | 9.457741 | - | 1.586111 | 0.30066 | 5 | -1247.02 | 2504.052 | 21.50947 | 1.06E-05 |
| -16.4964 | - | - | 1.871859 | 0.330384 | 4 | -1253.61 | 2515.243 | 32.69987 | 3.93E-08 |
| -11.1312 | 10.01418 | - | 1.304623 | - | 4 | -1254.02 | 2516.05 | 33.50674 | 2.63E-08 |
| -12.6494 | - | - | 1.566861 | - | 3 | -1260.9 | 2527.81 | 45.26669 | 7.34E-11 |
| -11.5005 | 5.156819 | 1.522347 | - | - | 4 | -1263.94 | 2535.884 | 53.34131 | 1.30E-12 |
| -13.595 | - | 1.854544 | - | - | 3 | -1265.03 | 2536.068 | 53.5254 | 1.18E-12 |
| -13.7149 | - | 1.897715 | - | -0.04475 | 4 | -1264.91 | 2537.827 | 55.28439 | 4.90E-13 |
| -11.5447 | 5.075363 | 1.532788 | - | -0.00597 | 5 | -1263.94 | 2537.896 | 55.3529 | 4.74E-13 |
| -1.87957 | 15.67414 | - | - | 0.164593 | 4 | -1271.51 | 2551.034 | 68.49064 | 6.65E-16 |
| -1.18695 | 15.57846 | - | - | - | 3 | -1273.33 | 2552.669 | 70.12636 | 2.93E-16 |
| -1.04461 | - | - | - | 0.171518 | 3 | -1285.01 | 2576.02 | 93.47675 | 2.50E-21 |
| -0.33416 | - | - | - | - | 2 | -1286.55 | 2577.096 | 94.55273 | 1.46E-21 |
| **T= 3-day post infection** |  |  |  |  |  |  |  |  |  |
| **-16.995022** | - | **2.03555664** | **0.55147102** | **-0.191400** | **5** | **-1274.6647** | **2559.35031** | **0** | **0.66632070** |
| -17.339435 | -1.5271051 | 2.103227925 | 0.542990627 | -0.196125 | 6 | -1274.4711 | 2560.971478 | 1.621168373 | 0.296245027 |
| -10.17935 | - | 1.612735042 | - | -0.251981 | 4 | -1279.5797 | 2567.173439 | 7.823129504 | 0.013332522 |
| -20.02456 | - | 2.158428152 | 0.760673763 | - | 4 | -1279.6290 | 2567.272004 | 7.921694212 | 0.012691392 |
| -10.768006 | -2.0079994 | 1.70988591 | - | -0.256854 | 5 | -1279.2650 | 2568.550983 | 9.200673352 | 0.006695492 |
| -20.133321 | -0.3961623 | 2.176817493 | 0.75977039 | - | 5 | -1279.6173 | 2569.255605 | 9.90529541 | 0.004707342 |
| -10.75746 | - | 1.5459523 | - | - | 3 | -1288.3825 | 2582.773391 | 23.42308112 | 5.46E-06 |
| -10.950153 | -0.64100947 | 1.57673391 | - | - | 4 | -1288.3559 | 2584.725752 | 25.37544172 | 2.06E-06 |
| 3.048364661 | 10.2602745 | - | -0.33813037 | -0.203665 | 5 | -1307.7367 | 2625.494291 | 66.14398116 | 2.89E-15 |
| 0.590754189 | 12.62890254 | - | - | -0.151574 | 4 | -1309.2742 | 2626.562503 | 67.21219267 | 1.69E-15 |
| -0.0436170 | 12.64090345 | - | - | - | 3 | -1311.3255 | 2628.659407 | 69.30909746 | 5.93E-16 |
| 0.963400705 | 11.57801632 | - | -0.15204647 | - | 4 | -1310.9919 | 2629.997816 | 70.64750614 | 3.04E-16 |
| 5.773130979 | - | - | -0.65828154 | -0.246481 | 4 | -1314.1221 | 2636.258252 | 76.90794214 | 1.33E-17 |
| 3.635925332 | - | - | -0.48025385 | - | 3 | -1318.5842 | 2643.176849 | 83.82653942 | 4.18E-19 |
| 1.231046881 | - | - | - | -0.143057 | 3 | -1320.6772 | 2647.362764 | 88.01245429 | 5.15E-20 |
| 0.637355231 | - | - | - | - | 2 | -1322.2186 | 2648.441465 | 89.09115502 | 3.00E-20 |

**Table S2.** Estimates for binomial generalised linear mixed effects models with variables that were retained following AIC selection. ‘Phage’ refers to initial phage inoculum size, whilst ‘log_phage’ and ‘log_cell’ indicate measured phage and cell densities, respectively, at each timepoint.

|  | Estimate | Std. Error | z value | Pr(>\|z\|) | 467 |
| --- | --- | --- | --- | --- | --- |
| **T= 1-day post infection** |  |  |  |  |  |
| (Intercept) | -31.4092 | 3.34962 | -9.37694 | 6.79E-21 |  |
| Phage | 0.364452 | 0.056302 | 6.473196 | 9.60E-11 |  |
| log_phage | 3.102939 | 0.357563 | 8.678031 | 4.03E-18 |  |
| **T= 2-day post infection** |  |  |  |  |  |
| (Intercept) | -22.6156 | 2.433671 | -9.29281 | 1.50E-20 |  |
| log_cell | 1.582378 | 0.225369 | 7.021265 | 2.20E-12 |  |
| log_phage | 1.391144 | 0.20461 | 6.799011 | 1.05E-11 |  |
| **T= 3-day post infection** |  |  |  |  |  |
| (Intercept) | -16.995 | 2.579733 | -6.58789 | 4.46E-11 |  |
| Phage | -0.1914 | 0.058546 | -3.26924 | 0.001078 |  |
| log_cell | 2.035555 | 0.220627 | 9.226229 | 2.80E-20 |  |

